# Dopamine dynamics in chronic pain: music-induced sex-dependent behavioral effects in mice

**DOI:** 10.1101/2024.03.05.583510

**Authors:** Montse Flores-García, África Flores de los Heros, Ester Aso, Jennifer Grau-Sánchez, Sebastià Videla, Antoni Rodríguez-Fornells, Jordi Bonaventura, Víctor Fernández-Dueñas

**Affiliations:** Pharmacology Unit, Department of Pathology and Experimental Therapeutics, School of Medicine and Health Sciences, Institute of Neurosciences, Universitat de Barcelona, Barcelona, Spain; Neuropharmacology & Pain Group, Neuroscience Program, IDIBELL-Bellvitge Institute for Biomedical Research, Barcelona, Spain; Research group on Complex Health Diagnoses and Interventions from Occupation and Care (OCCARE). University School of Nursing and Occupational Therapy of Terrassa, Autonomous University of Barcelona, Terrassa, Spain; Cognition and Brain Plasticity Unit, Department of Cognition, Development and Educational Psychology, Faculty of Psychology, University of Barcelona and Bellvitge Institute for Biomedical Research, Barcelona, Spain; Catalan Institution for Research and Advanced Studies (ICREA), Barcelona, Spain

**Keywords:** Music, chronic pain, anxiety, depression, dopamine, nucleus accumbens, fiber photometry

## Abstract

Chronic pain is a debilitating disease that is usually comorbid to anxiety and depression. Current treatment approaches primarily rely on analgesics, but they often neglect emotional aspects. Non-pharmacological interventions have been incorporated into clinics to provide a more comprehensive management of chronic pain. Among these interventions, listening to music is a well-accepted and cost-effective option. However, the underlying mechanisms of music-mediated pain relief remain insufficiently understood. Here, our aim was to evaluate the effects of music exposure in an animal model of chronic pain. First, we injected mice with the inflammatory agent complete Freund’s adjuvant (CFA) into the hind paw and housed them for 14 days with background music during their active period (Mozart K.205, overnight), or silence. The impact of music exposure on nociception and anxiety-like and depression-like behaviors was evaluated through different paradigms, including the hot plate, Von Frey, elevated plus maze, splash, and tail suspension tests. Additionally, we investigated whether music influences dopamine dynamics in the nucleus accumbens (NAcc), a pivotal region involved in pain processing, anhedonia, and reward. Our findings indicate that music exposure prevents the decreased NAcc activity observed in CFA-injected mice, linking with a sex-dependent reduction of allodynia, anxiety- and depression-like behaviors. Thus, females were more sensitive to music exposure. Collectively, our findings provide compelling evidence for the integration of music listening as a non-pharmacological intervention in chronic pain conditions. Moreover, the observed impact on the NAcc suggests its potential as a therapeutic target for addressing chronic pain and its associated symptoms.

## 2 Introduction

Chronic pain, with a global prevalence ranging from 10% to 40%, stands as a critical public health issue (Cohen, Vase, and Hooten 2021a; Goldberg and McGee 2011). It is not only a primary reason for seeking medical care but also exerts a substantial socioeconomic impact on society (Cohen, Vase, and Hooten 2021a; Goldberg and McGee 2011). Beyond its physical effects, chronic pain significantly impairs patients’ quality of life and mental well-being, often manifesting as comorbid symptoms like anxiety, anhedonia, and depression (Roughan et al. 2021; Cohen, Vase, and Hooten 2021b; Garland et al. 2020). Furthermore, emerging evidence suggests a reciprocal exacerbation between the physical and psychological symptoms, emphasizing the need for comprehensive multimodal approaches to managing chronic pain conditions (Roughan et al. 2021).

In recent years, there has been a paradigm shift in chronic pain management, acknowledging the need for a more holistic perspective to address this condition. Non-pharmacological interventions have emerged as valuable complements to analgesics to help individuals in better anticipating, perceiving, and responding to pain (Shi and Wu 2023). Music listening is a non-pharmacological intervention commonly accepted for its non-invasiveness, safety, cost-effectiveness, and user-friendly nature (Shi and Wu 2023; Lunde et al. 2019; Roy, Peretz, and Rainville 2008; Cepeda et al. 2006; Moss et al. 2023). Clinical studies assessing the impact of music listening on chronic pain consistently report enhanced pain relief, along with improvements in associated comorbidities such as anxiety and depression (Lunde et al. 2019; A Y R Kühlmann et al. 2018; Gutgsell et al. 2013; Lee 2016). Nevertheless, the neurological mechanisms underlying the effects of music listening in reducing pain and associated comorbidities remain mostly unknown. As we recently reviewed, the mesolimbic system, connecting the ventral tegmental area (VTA) and the nucleus accumbens (NAcc), has gained attention in the intersection of pain and mood disorders (Flores-García et al. 2023). Interestingly, listening to music has been shown to induce changes in neural activity, triggering the release of neurotransmitters such as dopamine or endogenous peptides in different brain areas, including the NAcc (Kuner and Kuner 2021; Martikainen et al. 2015; Moraes et al. 2018a; Mavridis 2015; Chanda and Levitin 2013; Ferreri et al. 2019; Blood and Zatorre 2001; Salimpoor et al. 2011). Additionally, music listening can induce alterations at the peripheral level (e.g., changes in heart rate), with the autonomic and descending pain-modulatory systems being the primary players in these changes (Lunde et al. 2019; Fu et al. 2023; Chanda and Levitin 2013).

The effects of music exposure in animal models, particularly in the context of pain and mood disorders, have gained significant attention. Numerous studies have investigated into how music interventions influence behavioral responses to pain, stress, and emotional states in different animal models (A. Y. Rosalie Kühlmann et al. 2018; Metcalf et al. 2019; Mao, Cai, and Lou 2022). It is important to note that animals may not listen to music in the same way humans do. For instance, human perception of music is complex and influenced by cultural, emotional, and cognitive factors (Lunde et al. 2019; Zatorre and Salimpoor 2013). Similarly, the human brain complexity allows for intricate processing of music, engaging different brain areas associated with emotions, prediction, memory, and reward (Lunde et al. 2019; Zatorre and Salimpoor 2013; Chanda and Levitin 2013). Nevertheless, animals demonstrate an ability to respond and discriminate rhythmic patterns, tones, and melodies, indicating that music can perceptibly affect different species (Ito et al. 2022; A. Y. Rosalie Kühlmann et al. 2018; Celma-Miralles and Toro 2020). For example, recent research has shown differences in animal responses when being exposed to forward and backward music, evidencing certain capacity to decode the internal structure of music (Xing et al. 2016). Besides, rodents display innate spontaneous beat synchronization and neural tuning in the auditory cortex (Ito et al. 2022). Despite these differences, animal studies could offer valuable insights into the potential benefits of music and its mechanisms of action. These insights could lead to the development of novel strategies to manage chronic pain conditions.

The ultimate goal of our investigation it to pave the way for more refined and effective treatment strategies, enhancing the quality of life for individuals suffering from chronic pain and its emotional comorbidities. Here, we used an animal model of chronic pain to contribute to the understanding of music-mediated effects in this condition. To achieve this, we first injected the inflammatory agent (complete Freund’s adjuvant, CFA) into the hind paw of mice (Spinieli et al. 2022; Fernández-Dueñas et al. 2007). Subsequent to CFA injection, animals developed sustained pain accompanied by anxiety- and depression-like behaviors (Spinieli et al. 2022; Hipólito et al. 2015; Schwartz et al. 2014a; Massaly et al. 2019). We assessed the influence of music exposure on nociceptive and emotional CFA-mediated effects, employing fiber photometry to investigate putative music-mediated modulation of dopamine dynamics in the NAcc.

## 3 Materials and Methods

### 3.1 Animals

Adult male and female CD-1 mice (animal facility of University of Barcelona) weighing 30-40 g were used. Animals were housed and tested in compliance with the guidelines provided by the Guide for the Care and Use of Laboratory Animals (Clark et al., 1997) and following the European Union directives (2010/63/EU). The University of Barcelona Committee on Animal Use and Care approved the protocol. Mice were randomly assigned to each experimental group and housed in standard cages with ad libitum access to food and water and maintained under a 12 h dark/light cycle (starting light period at 8:00 AM), 22°C temperature, and 66% humidity (standard conditions). All animal experimentation was carried out in a period comprehended between 9:00 AM to 6:00 PM by a researcher blind to treatments.

### 3.2 Reagents

Complete Freund’s adjuvant (CFA) was purchased from Sigma-Aldrich (St. Louis, MO, USA). The antibodies used were: rabbit anti-tyrosine hydroxylase antibody (1:1000, AB152, Merck Life Science SLU, Darmstadt, Germany), chicken anti-GFP (1:500, A10262, Thermo Fisher Scientific Inc, Waltham, Massachusetts, USA), donkey anti-rabbit Alexa Fluor 647 (1:1000, A31573, Thermo Fisher Scientific Inc) and goat anti-chicken Alexa Fluor 488 (Thermo Fisher Scientific Inc).

### 3.3 Experimental design

#### 3.3.1 Induction of chronic pain and behavioral assessment

Chronic pain was induced by the subplantar injection of 0.05 ml of CFA into the right hind paw, under brief anesthetic conditions induced with isofluorane (3 %), as previously reported (Fernández-Dueñas et al. 2007). Mice were randomly divided into two groups: vehicle- and CFA-injected. Of note, each group was subdivided into two groups depending on the sex (male, female). After 13 days the effects of CFA injection were evaluated on nociception, while the effects on anxiety-like and depression-like associated symptoms were assessed on day 14.

#### 3.3.2 Impact of music on chronic pain

CFA-injected mice were divided into two groups based on their housing conditions during the 14 days post-injection: 1) silence: ambient noise ≈ 40-45 dB, and 2) music exposure: Mozart K-205, 20:00 p.m. to 8:00 a.m., ≈ 55 ± 10 dB. Of note, each group was subdivided into two groups depending on the sex (male, female). After 13 days the effects of music were evaluated on nociception, while the effects on anxiety-like and depression-like associated symptoms were assessed on day 14.

#### 3.3.3 Effects of music on dopamine dynamics in the NAcc

Dopamine dynamics were assessed in the NAcc of female mice. First, we performed stereotaxic surgery to inject a dopamine biosensor (see below) and implant a fiber optic in the NAcc. After a 4-week period, we performed the subplantar injection of CFA and divided mice into two groups (silence, music exposure; *see* 3.3.2). Fiber photometry experiments were performed previous to CFA injection and 7 and 14 days later.

### 3.4 Evaluation of anti-nociceptive effects

#### 3.4.1 Hot plate test

Thermal nociceptive thresholds were evaluated using a hot plate apparatus (Ugo Basile instruments, Comerio, Italy), as previously described (Fernández-Dueñas et al. 2010). Briefly, mice were individually placed in the hot-plate metallic surface maintained at 48 ± 0.2 °C, and the latency of the injected hind paw withdrawal determined using a timer integrated into the system. A cut-off of 18 s was established to avoid tissue damage.

#### 3.4.2 Von Frey test

Mechanical nociceptive thresholds were evaluated using an automated von Frey-type dynamic plantar aesthesiometer (Ugo Basile). Mice were placed individually in transparent acrylic boxes with wire mesh floors and habituated for 1 h before testing. The tip of the von Frey filament was applied to the middle of the plantar surface of the hind paw with increasing force until the paw was withdrawn or a cut-off force of 50 g was reached. The force required to elicit a reflex withdrawal of the hind paw was automatically recorded. The lowest force required in three trials performed at 5 min intervals was considered to be the withdrawal threshold.

### 3.5 Evaluation of anxiety-like and depression-like behaviors

#### 3.5.1 Elevated plus maze test

The elevated plus maze (EPM) was used to assess anxiety-like symptoms (Walf and Frye 2007). In brief, mice were placed in the middle of the EPM apparatus, which consists of a cross composed of two open and two closed arms (16×5 cm) joined by a common central platform (5×5 cm), and let them to freely explore the maze. The total distance travelled and the time spent in both closed and open arms were calculated using a home-made Matlab script (The Mathworks Inc, Natick, MA, USA), which analyzed data after video analysis using the DeepLabcut (DLC) machine learning based package (Mathis et al. 2018). Briefly, a DLC project was trained to coordinately tracking various body parts (nose, right and left ear, center of the body and tail base). The training consisted of 200,000 iterations and 200 frames per video. The tracks obtained were analyzed using custom scripts in Matlab R2023b (The Mathworks Inc) to obtain the total distance travelled and the time spent in both closed and open arms. The percentage of time spent in the open arms was calculated as follows: (time in open arms / (time in open arms + time in closed arms))*100.

#### 3.5.2 Splash test

The splash test (Isingrini et al. 2010) was used to evaluate depression-like behavior (i.e. anhedonia). Each animal received on the dorsal coat two sprays from an atomizer spray containing a 10% sucrose solution. The viscosity of the sucrose solution induces grooming behavior in mice, serving as an index of self-care and motivational behavior (Isingrini et al. 2010). Mice were individually placed into round glass cylinders (20 cm in diameter) and the time spent grooming the splashed area was recorded for 5 min and scored by an experimenter blind to the treatment group.

#### 3.5.3 Tail suspension test

The tail suspension test (TST) was used to evaluate depression-like behavior (i.e. despair behavior) (Cryan, Mombereau, and Vassout 2005). The test consisted of suspending mice 50 cm above the floor by means of an adhesive tape, placed approximately 1 cm from the tip of the tail. Time of immobility was quantified for 5 min. Immobility was considered only when mice were hunging passively and motionless.

### 3.6 Fiber photometry

#### 3.6.1 Adeno-associated virus (AAV)

Adeno-associated viral particles encoding the dopamine biosensor Dlight1.3b (AAV-hSyn-Dlight1.3b; ETH Viral Vector Facility, University of Zurich, Switzerland) was used (Patriarchi et al. 2018).

#### 3.6.2 Virus delivery and optical fiber implantation

Animals were anesthetized using isoflurane in O_2_ (3% induction and 1.5% maintenance) and then placed in a stereotaxic apparatus (KOPF Instruments, Tujunga, CA, USA). The AAV preparation (500 nL) encoding Dlight1.3b (AAV1, titter 7,6 × 10^12 vg/mL) was slowly infused with a 2 µL Hamilton Syringe (Hamilton Company, Reno, NV, USA) at a rate of 50 nL/min unilaterally in the NAcc (relative to bregma: antero-posterior, 1.2 mm; medio-lateral, −0,7 mm; and dorsoventral, −4,2 mm. Next, an optical fiber (diameter 2.5 mm Ceramic Ferrule, 400 µm Core, 0.5 NA, 6 mm length) was implanted with the tip targeting 0.1 mm above the injection coordinates. Ceramic ferrules were secured to the skull using dental cement (Dentalon Plus, Kulzer Iberia S.A., Madrid, Spain) with the help of three skull-penetrating screws. Mice were removed from the stereotaxic instrument and placed over a heat pad in their home cages. Mice were then housed for four weeks to allow virus expression.

#### 3.6.3 Fiber photometry recordings

Fiber photometry recordings were conducted at three time points: a basal lecture 4 hours before CFA injection, and 7 and 14 days after CFA injection. Excitatory light was delivered using T-Cube™ LED drivers (Thorlabs, Ann Arbor, MI, USA) coupled with the integrated LEDs of Doric Minicubes (FMC4, Doric Lenses, Québec, QC, Canada). All signals were controlled and recorded using a Fiber Photometry Console (Doric) and Neuroscience Studio V6 (Doric). The emission light was optically filtered and registered using the built-in photodetector of the Minicubes and demodulated using the built-in algorithms of the Doric Neuroscience Studio. Changes in dopamine were recorded as changes in fluorescence emission at 530 nm excited by a 470 nm light source, while the same emission excited by 410 nm was used as isosbestic control. The signal was post-processed using Matlab custom-written codes. Briefly, the isosbestic signal was scaled to the Dlight1.3b signal using Iteratively Reweighted Least Squares fit. The biosensor signal (F) was decay -and artifact-corrected and normalized using the rescaled isosbestic signal (F_0_) using the formula: ΔFF= (F-F_0_)/F_0_. Peaks in the signal (dopamine transients) were detected and their properties (frequency, amplitude, and width) were calculated (Matlab) and statistically analyzed (GraphPad Prism 9.5.1, San Diego, CA, USA).

### 3.7 Immunofluorescence

At the end of the experiment mice were anesthetized and tissues fixed by the intracardial perfusion with 4% paraformaldehyde (PFA) diluted in phosphate-buffered saline (PBS). Brains were rapidly extracted and stored into 4% PFA solution for 24 hours and then placed into a 30% sucrose solution with 0,02% azide (in PBS). Brains were frozen and cut into coronal sections of 25 µm thickness by using a cryostat (Leica, Weltzar, Germany). The sections were stored at −20°C until use. Thereafter, sections were washed in PBS and permeabilized with 0,3% triton X-100 in PBS for 10 minutes. Sections were blocked with blocking buffer (0,05% triton X-100 + 2% bovine serum albumin in PBS) for one hour. Subsequently, they were incubated with the primary antibodies at 4 °C O/N, followed by the incubation with the secondary antibodies during 2 hours at room temperature. Sections were washed with 0,05% triton X-100 in PBS, incubated for 15 minutes with 1:1000 4′,6-diamidino-2-phenylindole (DAPI) solution, washed again with 0,05% triton X-100 in PBS and finally with PBS. The sections were mounted using a mowiol-based mounting medium with DABCO (1,4-Diazabicyclo[2.2.2]octane) anti-fading agent (Merck Life Science SLU). Images were acquired using a Carl Zeiss-Axio Imager.M2 microscope and processed with ZEN 3.9 software (Carl Zeiss Microscopy GmbH, Oberkochen, Germany).

### 3.8 Data analysis

Data are represented as mean ± standard error of mean (SEM) with statistical significance set at P < 0.05, unless indicated otherwise. The number of animals (n) in each experimental condition is indicated in the corresponding figure legend for each experimental condition. Comparisons among experimental groups were performed by two-way factor analysis of variance (ANOVA) followed by Tukey’s or Šídák’s multiple comparisons post-hoc test using GraphPad Prism 9.5.1 (San Diego, CA, USA), as indicated. Outliers were evaluated using the ROUT method assuming a Q value of 0.5% (GraphPad Prism 9.5).

## 4 Results

### 4.1 Evaluation of nociception, anxiety- and depression-like behaviors in chronic pain conditions

First, we aimed to validate our mouse model of chronic pain (Fernández-Dueñas et al. 2007), which consists of the unilateral injection of CFA into the hind paw, by evaluating nociception, anxiety- and depression-like behaviors (Fig. 1a). The subplantar injection of CFA induced an inflammatory response, which could be readily assessed measuring the paw thickness 1 day after injection and persisted for at least 14 days (Fig. 1b). Mice displayed hyperalgesia 13 days post-inflammation induction (Fig. 1c). We analyzed sex-based differences in the hot plate through two-way ANOVA, which revealed that there was an effect of inflammation [F_(1,44)_ = 29.03, P < 0.0001] but not an effect of sex [F_(1,44)_ = 0.36, P = 0.55], and that there was not an interaction between both factors [F_(1,44)_ = 0.68, P = 0.41]. Tukey’s post hoc analysis showed significant differences between both males and females following vehicle- or CFA-injection (P = 0.0123 and P = 0.0004, respectively). Similarly, in the Von Frey test we observed a diminution of the mechanical threshold in CFA-injected mice (Fig. 1d). The two-way ANOVA revealed that there was an effect of inflammation [F_(1,44)_ = 50.04, P < 0.0001] but not an effect of sex [F_(1,44)_ = 1.09, P = 0.30], and that there was not an interaction between both factors [F_(1,44)_ = 0.49, P = 0.48]. Tukey’s post hoc analysis showed significant differences between both males and females following vehicle- or CFA-injection (P < 0.0001 and P = 0.0006, respectively). These results further validated our previous results demonstrating that CFA-induced sustained hyperalgesia and allodynia over time (Fernández-Dueñas et al. 2007).

**Figure 1.**
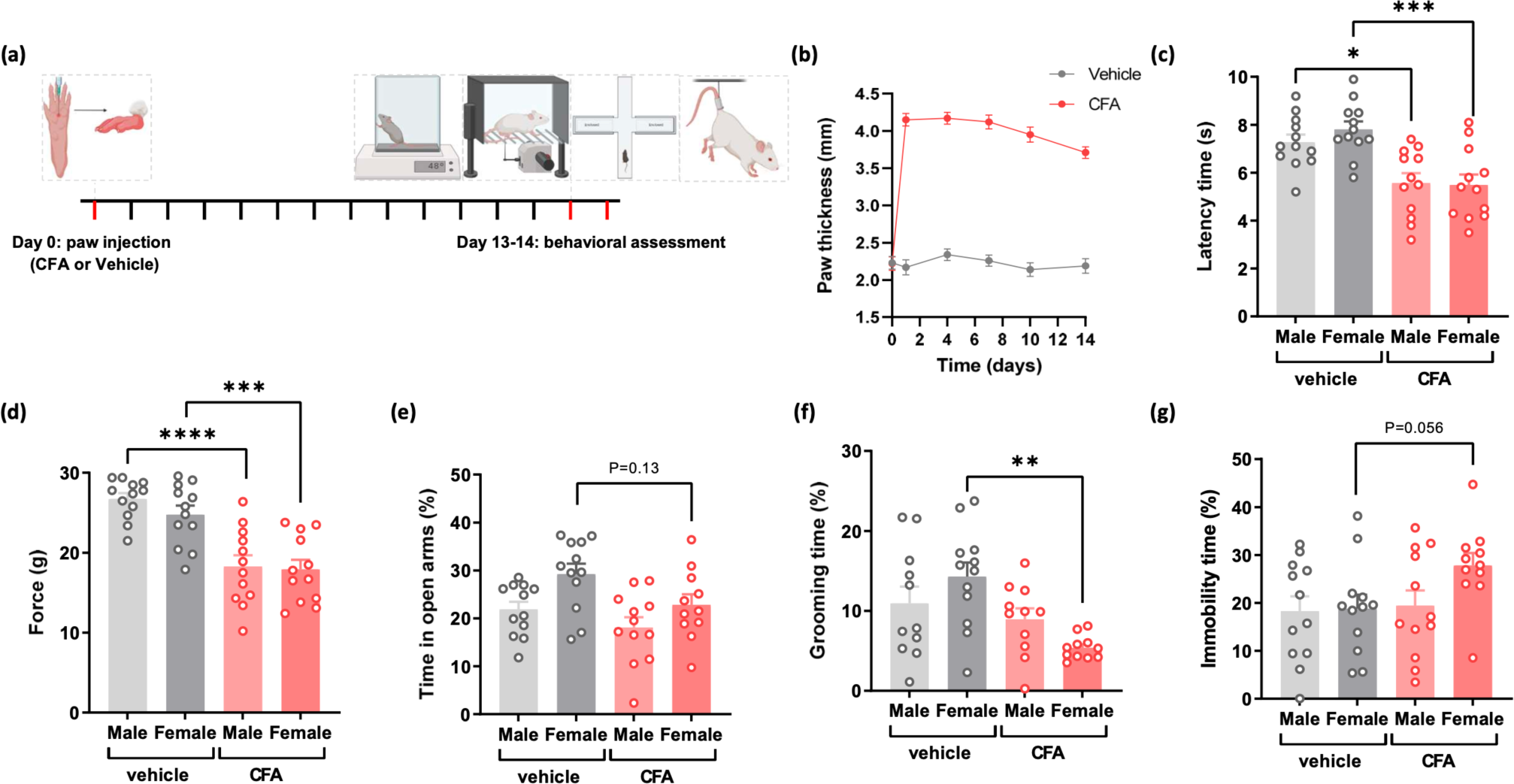
CFA-mediated nociception, anxiety- and depression-like behavior. **(a)** Experimental timeline: mice were subplantarly injected with CFA and the effects evaluated after 13 days using the hot plate and Von Frey tests, while anxiety-like and depression-like behaviors were evaluated after 14 days using the EPM, splash test and TST. **(b)** Effects of CFA injection on paw thickness. Paw thickness over time of vehicle- and CFA-injected mice (males and females). Data are presented as mean ± SEM of 24 animals/group. **(c)** Effects of CFA-injection in thermal hyperalgesia differentiated by sex. Latencies (s) compared between vehicle- and CFA-injected groups, including the sex. Data are presented as mean ± SEM of 12 animals/group. *P < 0.05, ***P < 0.001; Two-way ANOVA, Tukey’s post-hoc test. **(d)** Effects of CFA-injection in mechanical allodynia differentiated by sex. Force (g) inducing withdrawal compared between vehicle- and CFA-injected groups, including the sex. Data are presented as mean ± SEM of 12 animals/group. ***P < 0.001. ****P < 0.0001; Two-way ANOVA, Tukey’s post-hoc test. **(e)** Effects of CFA-injection in the percentage of time spent in open arms, compared between vehicle- and CFA-injected groups including the sex. Data are presented as mean ± SEM of 12 animals/group. P > 0.05; Two-way ANOVA, Tukey’s post-hoc test. **(f)** Effects of CFA injection on anhedonia. Time of grooming (%) of vehicle- and CFA-injected mice (males and females). Data are presented as mean ± SEM of 12 animals/group. **P < 0.01, Two-way ANOVA, Tukey’s post-hoc test. **(g)** Effects of CFA injection on despair behavior. Time of immobility (%) of vehicle- and CFA-injected mice (males and females). Data are presented as mean ± SEM of 12 animals/group. P > 0.05, Two-way ANOVA, Tukey’s post-hoc test.

Next, we aimed to explore whether CFA-injection leads to anxiety-like and depression-like behaviors. First, we examined the anxiety-like behavior associated to chronic pain using the EPM paradigm. Our data indicate that CFA-injected mice exhibited heightened anxiety-like behavior, spending less time in the open arms than vehicle-injected mice (Fig. 1e). The two-way ANOVA revealed that there was an effect of inflammation [F_(1,44)_ = 6.27, P = 0.0162] and sex [F_(1,44)_ = 8.84, P = 0.048], and that there was not an interaction between both factors [F_(1,44)_ = 0.44, P = 0.51]. However, Tukey’s post hoc analysis did not show significant differences between males or females following vehicle- or CFA-injection (P = 0.55 and P = 0.13, respectively). Next, we assessed whether CFA-injection also led to depression-like symptoms by using the splash test, a validated tool for assessing anhedonia (Isingrini et al. 2010). Fourteen days after the injection of CFA, a sucrose solution was sprayed on the animals back fur, and their behavior was recorded to score grooming time. Grooming time was lower in CFA-injected mice compared to vehicle-injected mice, indicating an anhedonic-like behavior (Fig. 1f). The two-way ANOVA revealed that there was an effect of inflammation [F_(1,44)_ = 12.11, P = 0.0012] but not an effect of sex [F_(1,44)_ = 0.007, P = 0.93], and that there was an interaction between both factors [F_(1,44)_ = 4.93, P = 0.03]. Tukey’s post hoc analysis only showed significant differences between females following vehicle- or CFA-injection (P = 0.011). To further explore another aspect of depression-like behavior we used the tail suspension test, which is indicative of despair behavior (Cryan, Mombereau, and Vassout 2005). CFA-injected mice exhibited increased immobility time compared to vehicle-injected mice (Fig. 1g). The two-way ANOVA revealed that there was an effect of inflammation [F_(1,44)_ = 8.48, P = 0.04] but not sex [F_(1,44)_ = 6.82, P = 0.07], and that there was not an interaction between both factors [F_(1,44)_ = 5.61, P = 0.10]. However, Tukey’s post hoc analysis did not show significant differences between males or females following vehicle- or CFA-injection (P = 0.99 and P = 0.056, respectively). Altogether, our findings provided evidence that CFA-injection induced anxiety-like and depression-like behaviors. These effects were particularly pronounced in female mice, with anhedonic-like behavior exhibiting the clearest and most significant effect.

### 4.2 Effects of music in nociception, anxiety-like and depression-like behaviors associated to chronic pain

Once validated our model of chronic pain, we investigated the impact of music exposure to prevent the observed CFA-mediated effects. In addition, we aimed to explore possible differences of music effects depending on sex. To this end, we injected CFA in the hind paw of different groups of male and female mice, and submitted them to a regimen of music exposure, or silence (*see* methods, Fig. 2a). First, we explored the potential antinociceptive effects of music. We examined possible differences between sexes, by assessing its antinociceptive effects in CFA-injected mice submitted or not to music exposure. We analyzed sex-based differences in thermal latencies through two-way ANOVA, which revealed that there was not an effect either of sex [F_(1,44)_ = 0.004, P = 0.95] or music exposure [F_(1,44)_ = 0.043, P = 0.84], and that there was not an interaction between both factors [F_(1,44)_ = 0.554, P = 0.46] (Fig. 2b). In contrast, music elevated the mechanical nociceptive thresholds. The statistical analyses revealed an effect of music exposure [F_(1,44)_ = 7.152, P = 0.01] but not sex [F_(1,44)_ = 0.649, P = 0.42], and that there was not an interaction between both factors [F_(1,44)_ = 1.788, P = 0.19]. Tukey’s post hoc analysis only showed significant differences between females following silence or music exposure (P = 0.0417), but not between males (P = 0.09). These findings indicated that music exposure reversed allodynia in females (Fig. 2c).

**Figure 2.**
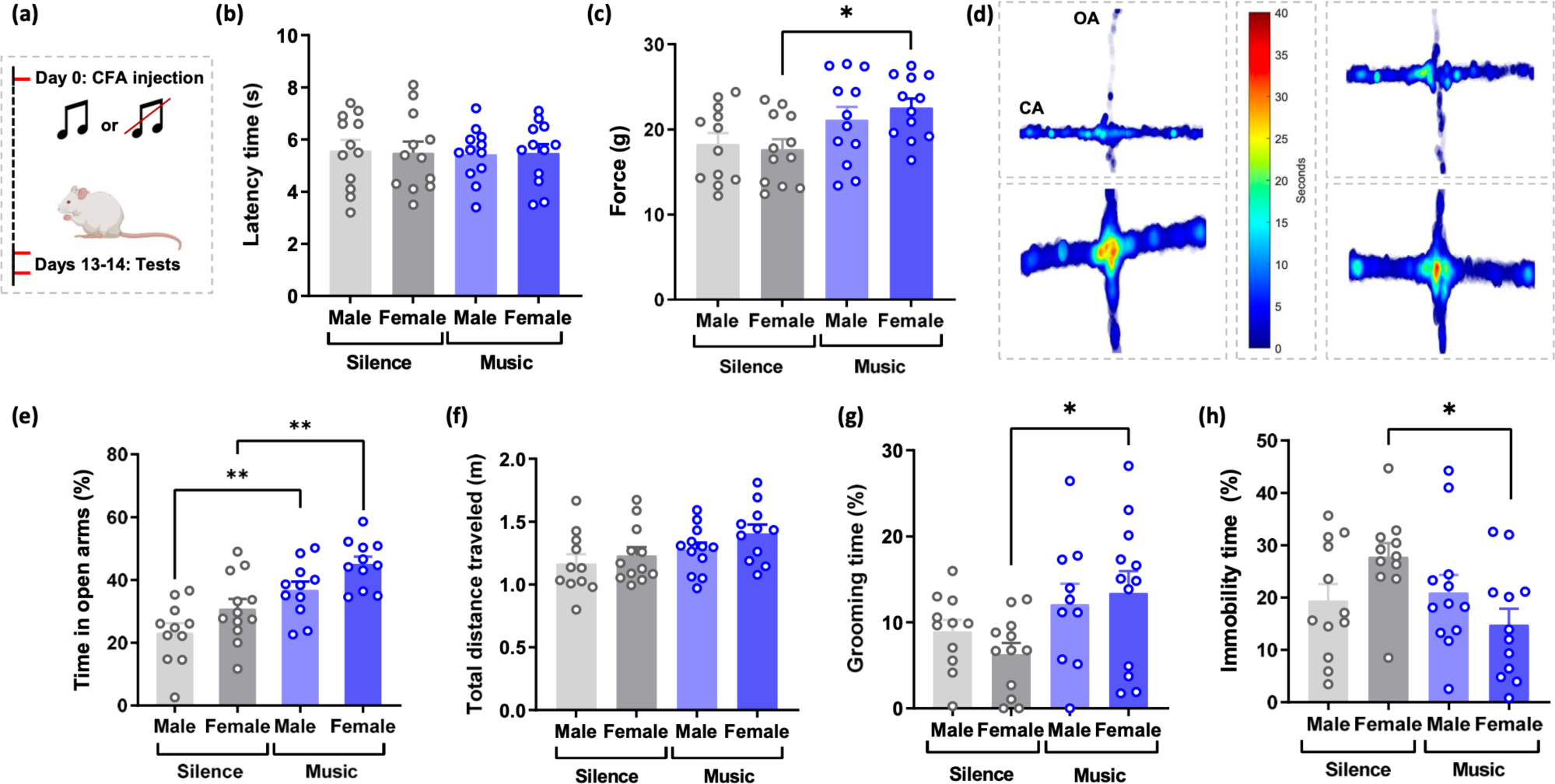
Effects of music on nociception, anxiety- and depression-like behaviors linked to chronic pain. **(a)** Experimental timeline: mice were subplantarly injected with CFA and the potential effects of music exposure (Mozart K-205, O/N) evaluated 13-14 days later. **(b)** Effects of music on CFA-induced thermal hyperalgesia differentiated by sex. Latencies (s) compared between silence and music groups, including the sex. Data are presented as mean ± SEM of 12 animals/group. P > 0.05; Two-way ANOVA, Tukey’s post-hoc test. **(c)** Impact of music exposure on CFA-induced mechanical allodynia differentiated by sex. Force (g) inducing withdrawal compared between silence and music groups, including the sex. Data are presented as mean ± SEM of 12 animals/group. *P < 0.05; Two-way ANOVA, Tukey’s post-hoc test. **(d)** Representative plots depicting the performance of CFA-injected mice: 1) male+silence, 2) female+silence, 3) male+music, 4) female+music, generated using the DLC software. The tracking of the animal across the maze is colored in relation to time. OA: open arm, CO: closed arm. **(e)** Quantification of the percentage of time spent in open arms, compared between silence and music groups including the sex. Data are presented as mean ± SEM of 12 animals/group. **P < 0.01; Two-way ANOVA, Tukey’s post-hoc test. **(f)** Distance travelled (m) compared between silence and music groups, including sex. Data are presented as mean ± SEM of 12 animals/group. P > 0.05; Two-way ANOVA, Tukey’s post-hoc test. **(g)** Effects of music on anhedonic behavior differentiated by sex. Grooming time (s) compared between silence and music groups including the sex. Data are presented as mean ± SEM of 12 animals/group. *P < 0.05; Two-way ANOVA, Tukey’s post-hoc test. **(h)** Impact of music on despair behavior differentiated by sex. Immobility time (s) compared between silence and music groups including the sex. Data are presented as mean ± SEM of 12 animals/group. *P < 0.05; Two-way ANOVA, Tukey’s post-hoc test.

Next, we evaluated the impact of music exposure, and possible differences between sexes in CFA-induced anxiety-like behavior. We used the machine learning based package DLC to analyze the video recordings and assess both the time spent in the arms and the distance travelled. In Fig. 2d are shown representative plots in which the time spent in the arms by the different groups is represented as a heat map. Quantification of these analyses showed that there was an increase in the percentage of time spent in the open arms of music-exposed mice compared to that of the silence group. Interestingly, this effect manifested both in male and female mice (Fig. 2e). The two-way ANOVA revealed that there was a significant effect of music exposure [F_(1,41)_ = 24.760, P < 0.0001] and sex [F_(1,41)_ = 8.195, P = 0.006], with no interaction between both factors [F_(1,41)_ = 0.906, P = 0.01]. Tukey’s post hoc analysis showed significant differences in both males and females when comparing silence vs music conditions (P = 0.008 and P = 0.004, respectively). Thus, data indicated that music exposure was effective in mitigating anxiety-like behavior across sex. We also assessed the total distance travelled during the test. Spontaneous activity may not be an optimal measure for anxiety-like behavior, but an increase in locomotor activity in the maze may also reflect anti-anxiety effects (Walf and Frye 2007). Indeed, music exposure augmented locomotor activity in music-treated mice but, in contrast to the percentage of time spent in the open arms, no sex differences were observed (Fig. 2f). The two-way ANOVA analysis revealed an effect of music exposure [F_(1,43)_ = 4.753, P = 0.04] but not sex [F_(1,43)_ = 2.078, P = 0.16], without interaction between the two factors [F_(1,43)_ = 0.558, P = 0.31]. Tukey’s post hoc analysis showed no significant differences between groups. Overall, our findings support that music effectively reduced anxiety-like behavior in CFA-injected mice.

After examining the impact of music on anxiety-like behavior associated to chronic pain, we assessed whether it was also effective at reducing depression-like symptoms. In the splash test, we observed that music exposure increased grooming time, thus reducing anhedonic-like behavior. The statistical analyses showed a main effect of music exposure [F_(1,41)_ = 6.988, P = 0.01] but not sex [F_(1,41)_ = 0.115, P = 0.74], with no interaction between both factors [F_(1,41)_ = 1.032, P = 0.32]. Tukey’s post hoc analysis showed significant differences in females under music exposure or silence (P = 0.049), but not in males (P = 0.7) (Fig. 2g). Similarly, we could observe an effect of music exposure when measuring despair behavior by means of the tail suspension test (Fig. 2h). The two-way ANOVA showed no main effects of music exposure [F_(1,43)_ = 3.427, P = 0.07] or sex [F_(1,43)_ = 0.132, P = 0.72], but revealed an interaction between both factors [F_(1,43)_ = 5.500, P = 0.02]. Tukey’s post hoc analysis indicated significant differences only in females with music exposure compared to females treated with silence (P = 0.0264). In summary, the effects of music on anhedonia and despair behavior were specifically observed in females.

### 4.3 Effects of music in dopamine release in the nucleus accumbens

Fiber photometry is a cutting-edge tool that allows real-time measurements of neurotransmitter release (Scimemi and Beato 2009; Patriarchi et al. 2020; 2018). The ability to quantify neurotransmitter release in a subsecond scale offers unprecedented opportunities to understand the neural mechanisms underlying different physiological and pathological processes (Scimemi and Beato 2009; Patriarchi et al. 2020; 2018). We hypothesized that the effects of music in chronic pain conditions might be mediated, at least in part, by alterations in dopamine dynamics in the NAcc (Flores-García et al. 2023). Accordingly, we investigated potential alterations in dopamine release over time following CFA injection and music exposure. Based on the previous behavioral results, we performed our experiments only in female mice. First, animals underwent stereotaxic surgery to inject the intensity-based fluorescent dopamine biosensor Dlight1.3b (Patriarchi et al. 2018) and implanting a fiber optic, and, after a 4-week period, we recorded extracellular dopamine transients in free moving animals (Fig. 3a). After CFA injection, we repeated photometry experiments at days 7 and 14 days of chronic inflammation with or without music exposure (Fig. 3a). Figure 3b illustrates the schematic representation of virus injection and insertion of a fiber optic into the NAcc to record spontaneous dopamine transients. Following the last fiber photometry recordings, mice were sacrificed, and the brains processed for immunohistochemistry to ensure virus expression in the NAcc and the correct optic fiber implantation (Fig. 3c).

**Figure 3.**
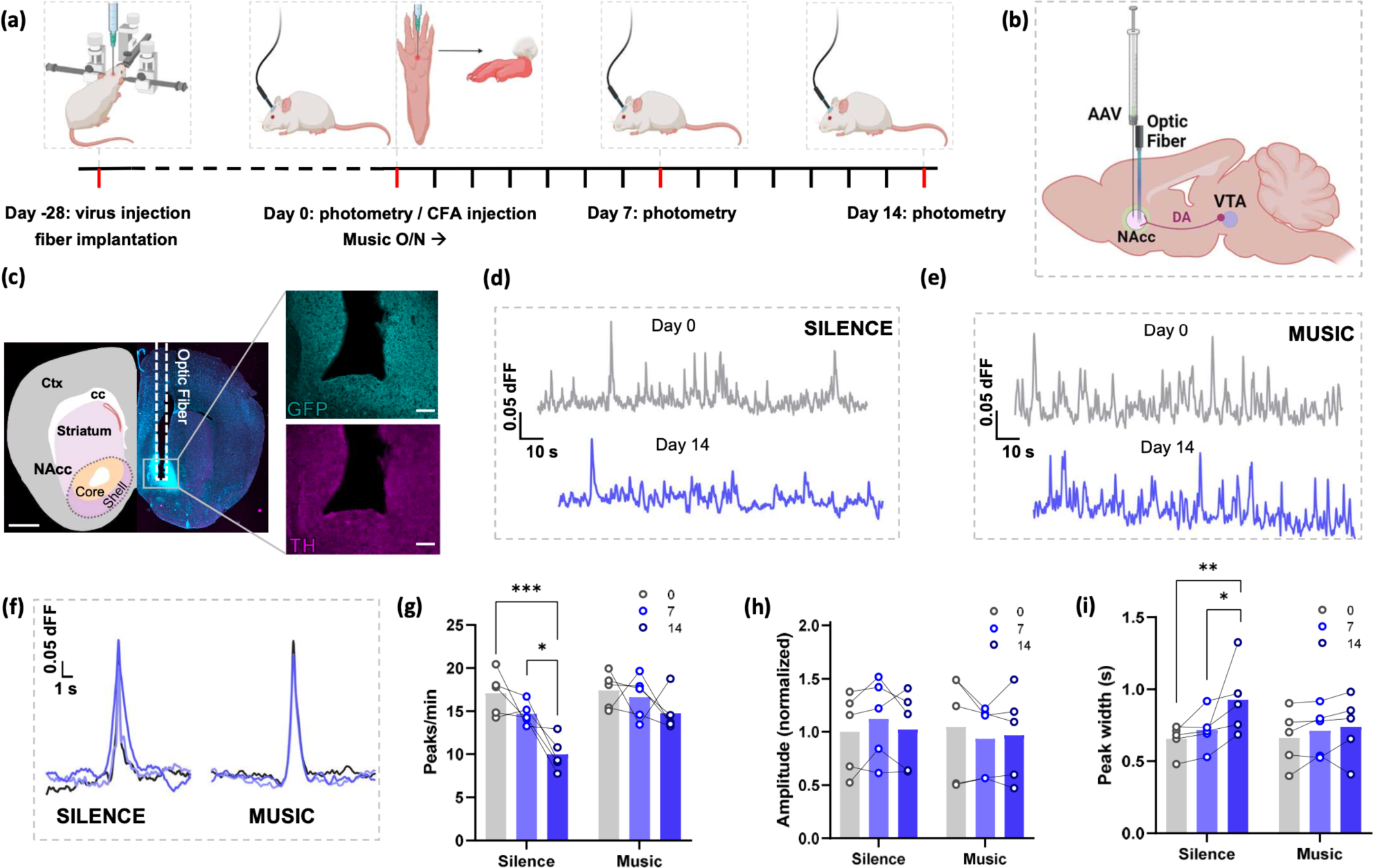
Assessment of dopamine dynamics in the nucleus accumbens. **(a)** Experimental timeline: mice were injected with a dopamine sensor virus followed by the insertion of a fiber optic into the NAcc, and 28 days later submitted to fiber photometry recordings. Next, mice were subplantarly injected with CFA and the potential effects of music treatment (Mozart K-205, O/N) in dopamine dynamics evaluated after 7 and 14 days. **(b)** Schematic illustration of surgical intervention for Dlight1.3b AAV injection and optic fiber implantation in the NAcc. **(c)** Whole brain slice showing the insertion of the optic fiber in the NAcc and immunofluorescence detection of the DLight1.3b sensor with an antibody targeted to the GFP moiety. Tyrosine hydroxylase (TH) and cell nuclei (with DAPI) were also immunolabelled. Scale bar = 1000 μm. In-set: magnified images (10x) of AAV expression (GFP, turquoise) and TH (magenta) in the vicinity of the optic fiber. Scale bar = 100 μm, respectively. A drawing of brain regions in the cerebral section overlaps on the left hemisphere. Ctx: cortex, cc: corpus callosum. **(d)** Representative traces showing spontaneous dopamine fluorescent transients at different timepoints (0 and 14 days) for the silence group. **(e)** Representative traces at different timepoints (0 and 14 days) for the music group. **(f)** Representative plot of overall peak alignment of the complete recordings at the different timepoints (0, 7, and 14 days) for both silence and music groups. **(g)** Effects of music treatment on spontaneous dopamine transients in CFA-injected mice. Number of peaks/min compared between silence and music groups (5 animals/group) at the different timepoints (0, 7, and 14 days). *P < 0.05, ***< 0.001; Two-way ANOVA, Šídák’s post-hoc test. **(h)** Peak amplitude, normalized to the mean of each condition at day 0, compared between silence and music groups (5 animals/group) at the different timepoints (0, 7, and 14 days). P > 0.05; Two-way ANOVA, Šídák’s post-hoc test. **(i)** Peak width compared between silence and music groups (5 animals/group) at the different timepoints (0, 7, and 14 days). *P < 0.05, **< 0.01; Two-way ANOVA, Šídák’s post-hoc test.

Photometry recordings were conducted in a home-cage environment, where animals had free exploration for 20 minutes. Animals previously underwent habituation for three days to habituate to the experimental conditions, including the tethering to the fiberoptic system. Representative traces from a non-music-exposed animal at timepoints 0 and 14 are presented (Fig. 3d), alongside traces from an animal exposed to music at the same intervals (Fig. 3e). Additionally, a representative alignment of dopamine transients average for each condition at different timepoints is illustrated (Fig. 3f). Quantitative analysis across animals showed a progressive decrease in the number of dopamine peaks following CFA injection, with a significant reduction observed at timepoint 14 (Fig. 3g). Importantly, music exposure exhibited a preventive effect on this decline. The two-way ANOVA revealed that there was a main effect of time [F_(2.16)_ = 12.180, P = 0.0006] and music exposure [F_(1,8)_ = 10.440, P = 0.012], without interaction between both factors [F_(2,16)_ = 2.488, P = 0.12]. Šídák’s post hoc analysis showed significant differences between timepoints 0 and 7 and timepoint 14 only in the silence group (P = 0.0008 and P = 0.0265, respectively). These results revealed a progressive reduction of spontaneous dopamine transients in the NAcc under chronic pain conditions. Music exposure prevented this decline, suggesting that it may exert a potential regulatory role in maintaining dopamine signaling dynamics in the context of chronic pain.

Peak analysis was conducted to discern potential alterations in dopamine transients induced by chronic pain and music exposure. Our results revealed no discernible effects of CFA-injection on peak amplitude, which remained consistent across all timepoints. Furthermore, music exposure did not exert a modulatory influence on peak amplitude (Fig. 3h). The two-way ANOVA revealed that there was not an effect of time [F_(2,16)_ = 0.211, P = 0.81] neither of music exposure [F_(1,8)_ = 0.798, P = 0.07], and that there was not an interaction between both factors [F_(2,16)_ = 2.462, P = 0.117]. Šídák’s post hoc analysis further substantiated that there were not differences between the timepoints (P > 0.05). On the other hand, CFA-injection led to an increase of peak width, an effect that music exposure prevented. The two-way within subject ANOVA revealed that there was an effect of time [F_(2,16)_ = 6.941, P = 0.007] but not music exposure [F_(1,8)_ = 0.342, P = 0.58], without interaction between both factors [F_(2,16)_ = 2.624, P = 0.10]. However, Šídák’s post hoc analysis disclosed significant differences between timepoints 0 and 7 and timepoint 14 exclusively in the non-music exposed group (P = 0.0058 and P = 0.0402, respectively). The widening of peaks suggests potential alterations in dopamine synaptic clearance, which music exposure prevented thereby maintaining the dopamine dynamics in the NAcc of CFA-injected mice.

## 5 Discussion

Our findings uncovered compelling sex-dependent effects of music exposure under chronic pain conditions. Music exposure did not modify thermal hyperalgesia but attenuated mechanical allodynia induced by CFA injection, an effect that was exclusively observed in females. This outcome aligns with an expanding body of literature supporting the efficacy of music across diverse rodent models (Metcalf et al. 2019; Mao, Cai, and Lou 2022; A. Y. Rosalie Kühlmann et al. 2018). Notably, recent research proposed that the effectiveness of music exposure in rodents would depend on the sound pressure level (that should be ≈ 50 dB) rather than consonancy (Zhou et al. 2022). Our music exposure regimen was set at ≈ 55 ± 10 dB, while control animals were housed in ambient noise conditions (≈ 40-45 dB). Thus, the possibility of a sound-dependent effect cannot be entirely ruled out. However, it is noteworthy that the observed behavioral effects align with prior studies investigating similar music regimens and sound levels (Li et al. 2010). Importantly, some of these studies demonstrated that employing non-consonant or unpleasant music at comparable sound pressure levels led to the loss of music-mediated effects (Tavakoli et al. 2012; Moraes et al. 2018b; A. Y. Rosalie Kühlmann et al. 2018).

It is important to note that we only observed music-dependent antinociceptive effects in females. These sex-specific effects are in accordance with numerous reports highlighting variations in pain perception and analgesic responses between male and female individuals (Bartley and Fillingim 2013; Nater et al. 2006). Hence, considering sex as a critical factor in chronic pain research appears imperative. Although our study did not investigate into the potential mechanisms explaining the distinct antinociceptive effects of music exposure, existing research has extensively explored changes in hormonal modulation (Mogil 2012; Manolagas and Kousteni 2001; Craft 2007; Kowalczyk et al. 2006). These changes would make females more susceptible to interventions that could modulate neuroendocrine responses, such as music (Chanda and Levitin 2013). On the other hand, it has been proposed that estrogens may influence the excitability of nociceptive neurons, alter neurotransmitter’s release, or interact with the endogenous opioid system (Xu, Nan, and Lan 2020; Fitzgerald et al. 2023; Huhn, Berry, and Dunn 2018). For instance, estrogens can influence the expression and function of opioid receptors (Chai et al. 2017) potentially enhancing the sensitivity of females to interventions modulating the opioid system, such as music (Mas-Herrero et al. 2023; Zatorre and Salimpoor 2013).

We explored the impact of music exposure in anxiety- and depression-like behavior associated with CFA-induced pain. In both male and female mice, music exposure exerted anxiolytic effects, aligning with previous studies highlighting the impact of music on anxiety (Bradt, Dileo, and Shim 2013; Fu et al. 2023). These studies investigated into the alterations induced by music on various systems, including the immune and autonomic systems, both recognized for their sensitivity to music interventions (Rebecchini 2021). On the other hand, we also observed an effect of music exposure on depression-like symptoms, specifically in females. Interestingly, our findings are in accordance with previous studies demonstrating the antidepressant effects of music (Fu et al. 2023; Papadakakis, Sidiropoulou, and Panagis 2019). In these investigations, music exposure was shown to restore homeostasis in the hypothalamus-pituitary-adrenal axis, prevent oxidative stress, and counteract neurotrophic factor deficits (Fu et al. 2023; A. Y. Rosalie Kühlmann et al. 2018). However, these studies did not specifically explore the influence of sex. The mechanisms through which music may influence depression-like behavior, particularly in females, are insufficiently understood. Taken together, our results highlight the importance for accounting for sex-specific factors in the therapeutic use of music for managing anxiety and depression associated to chronic pain.

Possible mechanisms explaining the benefits of music listening on chronic pain and associated symptoms may involve the modulation of various neurotransmitter systems, including the serotoninergic and dopaminergic pathways (Papadakakis, Sidiropoulou, and Panagis 2019; Mavridis 2015; Moraes et al. 2018a; Martikainen et al. 2015; Ren et al. 2021). In our recent review, we highlighted the convergence of mechanisms in chronic pain and depression, pointing the ventral tegmental area (VTA) as a potential therapeutical hub (Flores-García et al. 2023). Chronic pain conditions are often associated with alterations in the mesolimbic pathway, in which dopaminergic VTA neurons project to the NAcc (Ren et al. 2021; Kuner and Kuner 2021; Tanasescu et al. 2016). Simultaneously, the mesolimbic pathway is involved in the regulation of both mood and reward, and alterations in this pathway have been identified in the context of depression conditions (Nestler and Carlezon 2006; Heshmati and Russo 2015; Shirayama and Chaki 2006). We took advantage of the Dlight1.3b biosensor, which provided high temporal resolution (sub-seconds scale) to monitor the spatiotemporal dynamics of dopamine signals in the NAcc (Patriarchi et al. 2018). It is worth mentioning that the Dlight series of biosensors exhibit faster off-kinetics compared to other indicators like GRAB_D2_, hence allowing to detect changes in phasic dopamine release, while GRAB_D2_ would be more adequate to measure tonic dopamine (Patriarchi et al., 2020; Sun et al., 2020).

We conducted recordings of spontaneous dopamine transients in freely moving animals under basal conditions and following both CFA injection and music exposure. Our data demonstrate that music exposure effectively prevented the reduction in the frequency of spontaneous dopamine transients induced by chronic pain. This observation suggests that chronic pain may induce neuronal changes, potentially affecting the firing patterns or the excitability of dopaminergic neurons (Gee et al. 2020; Schwartz et al. 2014b). Among the potential causes for this phenomenon is the impairment of synaptic dopamine transmission, where alterations in dopamine reuptake or vesicular release could influence the occurrence of spontaneous transients (Schwartz et al. 2014b; Scimemi and Beato 2009). Moreover, we observed that the width of the peaks was wider, indicating potential impairment of dopamine reuptake and/or degradation, likely linked to changes in the kinetics of synapse clearance (Schwartz et al. 2014b; Scimemi and Beato 2009). Under normal conditions, dopamine diffusion is limited by various factors, particularly the dopamine transporter (DAT). Impairment of these mechanisms of synapse clearance could enhance the diffusional spread of dopamine from its point of release (Silm et al. 2019; Ford 2014). Interestingly, the peak amplitude remained unchanged, contrasting with findings in other studies (Wood et al. 2007; Xie et al. 2014; Gee et al. 2020), although the lack of effect on peak amplitude might be due to the biosensor sensitivity and expression. Potential disparities in these results could also originate from differences in the models used (acute vs. chronic pain) or the sex (male vs. female) of the animals (Gee et al. 2020; Wood et al. 2007; Xie et al. 2014). Therefore, the changes induced by CFA injection seemed to influence the occurrence rather than the intensity of spontaneous dopamine transients. The sustained decrease in dopamine levels may lead to a reduction in overall activity within the NAcc (Salinas et al., 2023; Schwartz et al., 2014). Collectively, it appears evident that chronic pain induces alterations in the VTA dopaminergic neurons projecting to the NAcc, and music exposure prevents such alterations. These findings agree with previous studies indicating that dopamine release increases in the mesolimbic system following music exposure (Blum et al. 2017; Menon and Levitin 2005; Ferreri et al. 2019; Blood and Zatorre 2001; Salimpoor et al. 2011).

In conclusion, our findings indicate that music exposure prevented the neurobiological adaptations observed in chronic pain, linking with antinociceptive effects and a reduction of anxiety-like and depression-like behaviors. Our findings not only provide support for the integration of music listening as a non-pharmacological intervention for a more comprehensive management of chronic pain, but also contribute to elucidating the underlying mechanisms of music-induced effects in chronic pain conditions.

## 7 Author Contributions

Substantial contributions to the conception or design of the work (A.R.-F., J.B., V.F.-D.); or the acquisition (M.F.-G., A.F.-H., E.A., V.F.-D.), analysis (M.F.-G., J.B., V.F.-D.), or interpretation of data for the work (all authors). Drafting the work or revising it critically for important intellectual content (all authors). Final approval of the version to be published (all authors). Agree to be accountable for all aspects of the work in ensuring that questions related to the accuracy or integrity of any part of the work are appropriately investigated and resolved (all authors).

## 9 Acknowledgments

We thank Esther Castaño and Benjamín Torrejón from the Scientific and Technical Services (SCT) group at the Bellvitge Campus of the University of Barcelona for their technical assistance. We also thank Koosje De Ru, Master student from University Utrecht for her help on the cerebral section drawing.

